# Neural coding in gustatory cortex reflects consumption decisions: Evidence from conditioned taste aversion

**DOI:** 10.1101/2024.02.28.582584

**Authors:** Martin A. Raymond, Ian F. Chapman, Stephanie M. Staszko, Max L. Fletcher, John D. Boughter

## Abstract

Taste-responsive neurons in the gustatory cortex (GC) have been shown to encode multiple properties of stimuli, including whether they are palatable or not. Previous studies have suggested that a form of taste-involved learning, conditioned taste aversion (CTA), may alter the cortical representation of taste stimuli in a number of ways. We used miniscopes to image taste responses from a large population of neurons in the gustatory cortex of mice before and after CTA to NaCl, comparing taste responses in control and conditioned mice. Following conditioning, no significant effects on the number of responsive cells, or the magnitude of response to either NaCl or other taste stimuli were found. However, population-level analyses showed that in mice receiving a CTA, the representation of NaCl diverged from other appetitive stimuli in neural space and moved closer to that of aversive quinine. We also tracked extinction of the CTA in a subset of animals and showed that as NaCl became less aversive, the neural pattern reverted to match the behavior. These data suggest that the predominant function of the taste representation in GC is palatability; the neuronal response pattern to stimuli at the population level reflects the decision of the animal to consume or not consume the stimulus, regardless of quality or chemical identity.

## Introduction

The gustatory cortical area (GC), located within insular cortex in mammals, plays an important role in taste-related learning and feeding decisions (Oliveira-Maia et al., 2012; Yiannakas and Rosenblum, 2017; Boughter and Fletcher, 2021). While electrophysiological studies with rats and mice show that taste quality and stimulus concentration can be encoded in the activity of GC neurons, much evidence has accrued that ensemble and population activity is also organized relative to the hedonic character of taste stimuli, ultimately reflecting the decision of the animal to consume or not consume (Stapleton et al., 2006; Jezzini et al., 2013; Li et al., 2016; Sadacca et al., 2016; Fletcher et al., 2017; Levitan et al., 2019; Bouaichi and Vincis, 2020; Chen et al., 2021). Recently, we used miniscope imaging of GC in mice to demonstrate that repeated familiarity with taste stimuli resulted in an increase in correlated activity over days among stimuli that were consumed in a similar fashion in lick tests (Staszko et al., 2022). Other forms of taste- or ingestive-based learning have also been shown to modify the cortical representation of taste stimuli, most notably conditioned taste aversion (CTA) learning (Yamamoto et al., 1989; Accolla and Carleton, 2008; Moran and Katz, 2014; Arieli et al., 2022).

CTA learning occurs when neutral or appetitive tastes are temporally associated with illness, with the result that animals will display subsequent aversion to those tastes. A large body of research has characterized potential neural circuits and mechanisms that underlie CTA formation and expression, including evidence for a significant role of GC (Braun et al., 1982; Yamamoto et al., 1995; Grossman et al., 2008; Barki-Harrington et al., 2009; Schier et al., 2014; Lavi et al., 2018; Kayyal et al., 2019; Abe et al., 2020; Yiannakas et al., 2021; Jung et al., 2022; Kolatt Chandran et al., 2023). What is less certain is whether and how CTA affects the neuronal activity of the GC itself, and whether a change in activity following learning directly reflects or predicts aversion of the conditioned taste stimulus (CS). Prior research suggests that the effects of CTA on neural activity in the GC takes the form of two conceptually distinct – though potentially synergistic – phenomena: Enhancement of the response to the conditioned stimulus (CS), and/or a shift in the neural representation of the CS reflecting its altered palatability. The former possibility may point to an increase in CS salience (Yasoshima and Yamamoto, 1998; Wilkins and Bernstein, 2006; Lavi et al., 2018). Other studies have reported changes in neuronal representation in the GC that indicate a shifting of the hedonic value of the CS from neutral or appetitive towards aversion (Accolla and Carleton, 2008; Moran and Katz, 2014; Lavi et al., 2018; Arieli et al., 2022).

Describing the neural representations of tastes in terms of palatability is not uncomplicated. While taste stimuli are often described as categorically palatable or unpalatable, the behavioral response of an animal to a given taste stimulus is subject to variability as a function of concentration, mixture, the simultaneous or even expected availability of other taste stimuli, and the animal’s physiological state and prior experience. For our purposes, we consider palatability a descriptive rather than predictive phenomenon, determined strictly by the behavioral response of an animal to a given taste stimulus; a stimulus is palatable to the extent that an animal is willing to consume it. The disconnect between neural imaging or recording and naturalistic consummatory behavior often fails to capture the complexity of taste experience, leaving a gap in our understanding of the how the cortical representation of taste is altered following learning. In the current study, we conducted calcium imaging of neurons in GC in freely moving mice via head-mounted fluorescent miniscopes, during licking behavior before and after CTA. This allowed us to observe activity in GC in real time over the course of learning, and to do so with minimal interference in the behavior of the animal.

## Methods

### Animals

Adult male (n=8) and female (n=7) C57BL/6J mice (The Jackson Laboratory, Bar Harbor, ME) were used for all miniscope experiments. The mean age was 180 days at the start of behavioral testing, with mean body weights of 28.7 g (males) and 22.6 (females). Animals were maintained on a standard 12-hour light/dark cycle and were group-housed in standard plastic shoebox cages (28 × 17.5 × 13 cm) with *ad libitum* chow and water. After lens implantation, mice were moved to individual housing to avoid damage to the lens imaging surface. All procedures were approved by the University of Tennessee Health Science Center Institutional Care and Use Committee.

### Surgical Procedure

Mice (mean age 108 days) were anesthetized using isoflurane (4-5% induction, 1-2% maintenance) and secured in a stereotaxic apparatus (David Kopf Instruments). Carprofen (5.0mg/kg) and dexamethasone (0.2mg/kg) were administered subcutaneously proceeding surgery. The scalp was then depilated and sanitized, an incision was made from Bregma to Lambda, and a craniotomy was performed above gustatory cortex (anterior +1.1 mm, lateral 3.3mm relative to Bregma) using a dental drill (Osada Inc). A micropipette was lowered into the craniotomy window to a depth of 1.75 mm relative to the brain surface, the tissue was allowed to settle for 10min, and 500 nL of AAV1.Syn.GCaMP6s.WPRE.SV40 (Addgene) were injected at a speed of 15 ul/sec using a Nanoject II (Drummond Scientific). The micropipette was removed 10 minutes after virus injection, at which point a gradient refractive index (GRIN) lens was implanted. GRIN lenses (4.1mm long by 1mm diameter; Inscopix) were stereotaxically implanted into insular cortex (e.g., Figure 2A-B) via the same craniotomy window to a ventral depth of 1.75 mm relative to the brain surface using a custom holder. The lens was first secured using cyanoacrylate glue, and then the entire skull was covered with dental cement (Ortho-Jet, Lang Dental) to seal the surgical area and act as a stable platform for imaging. The surface of the lens was covered with a custom 3d-printed cap, which was glued in place with cyanoacrylate to prevent damage prior to the baseplate procedure. Animals were given antibiotic food (Uniprim, Fisher Scientific) and a daily subcutaneous injection of carprofen/dexamethasone for five days following surgery. Animals were then given approximately 6-8 weeks recovery, both to allow for viral expression, and to facilitate optimal healing and clearing of the imaging window (Staszko et al., 2022). To attach baseplates, animals were anesthetized with isoflurane and placed back in the stereotaxic apparatus. A custom 3D-printed holder (adapted from ONE Core) mounted to the stereotaxic was used to precisely position a miniscope over the lens. The baseplate was then attached on the headcap using dental cement, along with a custom lightweight metal head bar, affixed with cyanoacrylate.

### Lickometer Acclimation and Training

All behavioral procedures were conducted using a contact lickometer which includes a test chamber, a shutter-controlled access port, and the ability to present up to 16 stimulus bottles (Davis MS-160, DiLog Instruments). The basic features and operation of this device have been previously described (St John and Boughter, 2009; Staszko et al., 2022). The number and timing of licks to each stimulus bottle were recorded in the lickometer software and by a custom microcontroller peripheral. Stimuli were presented at room temperature. Mice were acclimated to the lickometer and trained to lick filtered water over the course of 6 days: On day 1, water was removed from the home cage of the mice (all daily fluid intake was delivered in the lickometer for the duration of behavioral testing), and animals were placed in the test chamber for 10 minutes, with no stimulus presentations. On days 2 and 3, mice were given 10 min of free access to water in the lickometer. On days 4 and 5, animals were given 30 trials of water, with a trial duration of 5 sec and an inter-trial-interval (ITI) of 7.5 sec. On day 6, animals were again given 30, 5 sec trials of water, with an extended ITI of 60 sec.

### Taste Exposure and Conditioned Taste Aversion

Following training, animals were tested for 2 days with a panel of multiple tastes and water (pretest 1 and 2). Stimuli included 0.5 M sucrose, 0.3 M sodium chloride (NaCl), 0.02 M citric acid, 0.01 M quinine hydrochloride (high QHCl), 0.03 mM quinine hydrochloride (low QHCl), and 0.3 M potassium chloride (KCl). Concentrations were chosen based in part on our previous studies (Fletcher et al., 2017; Staszko et al., 2022). For each test session, the multi-taste panel consisted of 5 sec trials of each stimulus, alternating with 5 sec trials of water with an ITI of 60 seconds. The stimuli were presented in pseudorandom order in 2 blocks, constituting a total of 12 stimulus trials and 12 water rinses. On the day following the second taste panel presentation, animals were given access in the lickometer (conditioning session) to either 0.2 M lithium chloride (LiCl; CTA group, N = 8) or 0.2 M NaCl (control group, N = 7). Although these salts have similar orosensory properties to mice, ingestion of LiCl (but not NaCl) causes gastric malaise, which leads to subsequent avoidance of equimolar NaCl (Glatt et al., 2016). Concern over latent inhibitory effects on conditioning (0.3 M NaCl was included in the pre-test stimulus panel) was mitigated by the relatively brief and limited pre-exposure to NaCl (Lubow, 2009).

Access during the conditioning session was limited to a single presentation lasting 20 min or 1000 licks, whichever elapsed first. In this way, we attempted to minimize potential effects of meal size, as mice tested in this way will commonly consume about 3 times as much NaCl vs LiCl (Glatt et al., 2016). Following conditioning, mice were given a “rest day” with 20 min free access to water in the Davis Rig. On the following day, mice were tested (post-test) with the same taste panel from days 1-2 to assess the impact of conditioned taste aversion. The sequence of training and testing sessions is visualized in Figure 1A.

**Figure 1.**
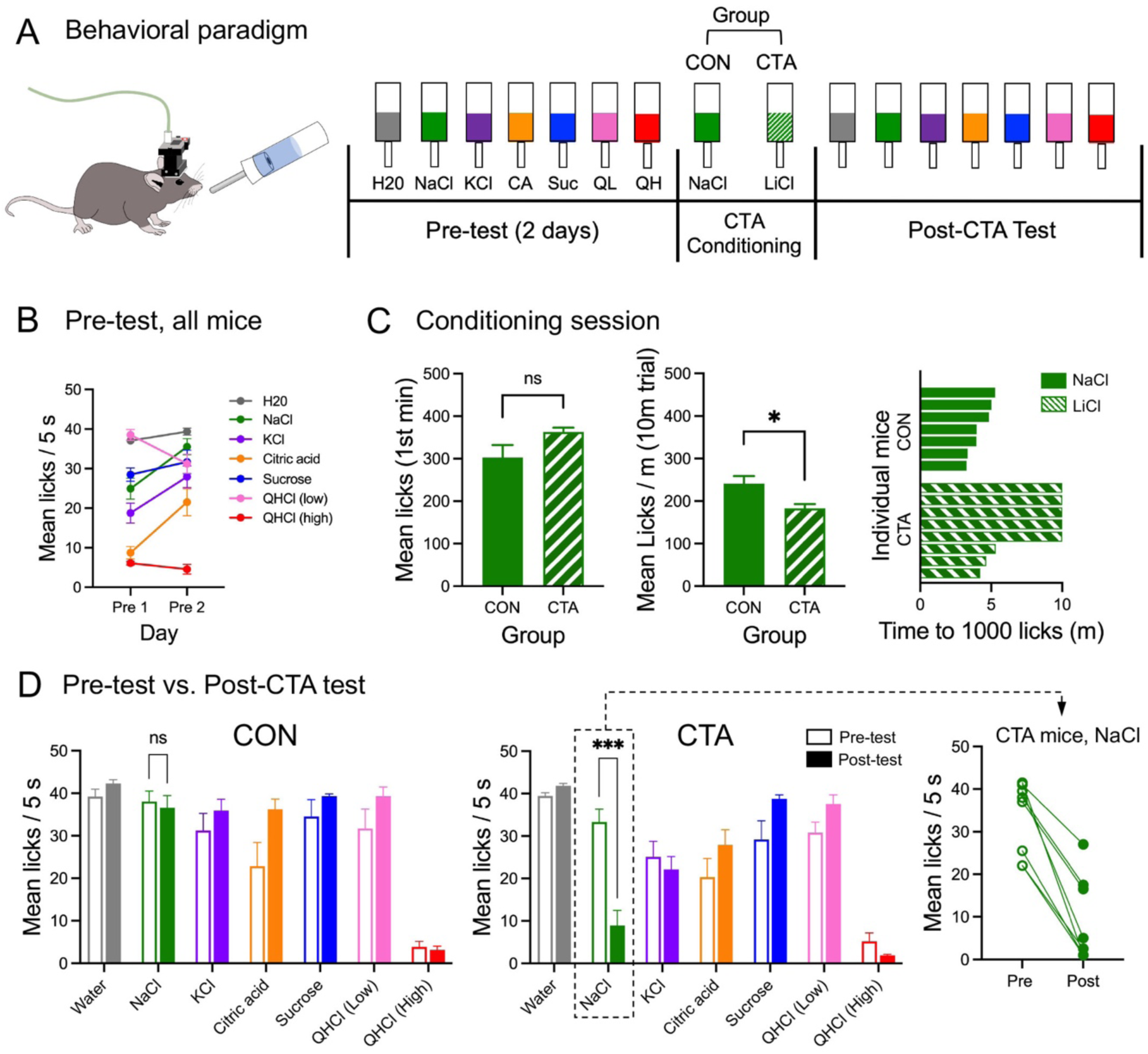
Behavioral testing: paradigm and results. A. Freely-moving mice with head-mounted miniscopes were tested with a short-trial battery of taste stimuli for 2 days (pretest); on conditioning day, they were given a single access period with either 0.2 M NaCl (control group; CON) or 0.2 M LiCl (CTA group). This was followed by a post-test day with the taste battery. B. Average licks per trial (all mice combined) recorded to each stimulus over the 2 pre-test days. C. Drinking behavior during conditioning; mean licks of the stimulus in the first minute, mean licks per minute averaged over the 10-min trial and for each mouse the time (min) elapsed before reaching 1000 licks. D. mean licks per trial for each stimulus, before (pre-test day 2) and after (post-test) for control and CTA animals, respectively. Inset: NaCl licking in the post-test was reduced for all CTA mice indicating successful generalization of the aversion. Asterisks (panels C, D) indicate significance in t-tests or post-hoc tests.

### Extinction of Conditioned Taste Aversion

Following the post-test, the testing paradigm was altered so as to facilitate extinction of the conditioned aversion. All mice were given a second post-test day, where they were presented with a modified taste panel in which the first 12 taste presentations were all 0.3 M NaCl. This was followed by a block of randomized taste presentations as in the previous sessions; each taste was presented once in a randomized series interleaved with water trials, for an additional 12 presentations. Given the highly divergent context of the 12 consecutive NaCl presentations relative to the single NaCl trial in the randomized series, these were treated as two different stimuli in analysis: the 12 consecutive trials were classified as “NaCl-Forced”. As in the previous phase of the experiment, all stimulus presentations were set to a 5-sec duration, with a 60 sec ITI. Presenting a series of initial NaCl trials drove high consumption of NaCl during this series, facilitating extinction of the conditioned aversion. This procedure was then repeated on a subset of the CTA mice (N = 4) for an additional 2 days, by which point the consumption of NaCl in the 2^nd^ randomized block had recovered to pre-CTA levels.

### Miniscope Imaging

Imaging was conducted using UCLA V4 miniscopes (www.miniscope.org; Cai et al., 2016). Tangling of the miniscope’s coaxial cable during freely-moving behavior was prevented by a custom-built commutator (ONE Core). Our previous study demonstrated that the presence of the miniscopes themselves did not alter brief-access taste behavior relative to non-miniscope wearing controls (Staszko et al., 2022). All videos were recorded at 30 FPS using the UCLA miniscope acquisition software. Simultaneous video recording of animal behavior in the chamber was collected using a webcam (Logitech) mounted above the Davis Rig. Gain, exposure, and LED intensity varied slightly between animals but were held constant across imaging days. All video segments collected in a single session were concatenated in Fiji, and imaging sessions were spatially down-sampled to half of their original area (a factor of 0.71 linearly), and temporally to 10 FPS to increase computational efficiency. Further pre-processing of imaging data, including motion correction and trace extraction, was conducted using the Python (3.6) implementation of CaImAn (Giovannucci et al., 2019). Rigid motion correction was used. Minimum spatial correlation and fluorescence peak-noise-ratio values for constrained non-negative matrix factorization with endoscopes (CNMFe) were determined by individual based on summary images. After running the CNMFe algorithm, outputs were further filtered by applying a minimum signal-to-noise ratio of 15.0, meaning the fluorescence signal had to reach a value of at least 15x the determined “noise” value for a given ROI to be accepted as a component. Accepted and rejected components were evaluated manually, and CNMF threshold values were adjusted if necessary, until false positives and negatives were minimized based on data visualization, on a per-animal basis. Threshold values, once established, were kept constant for all days. Identified components are categorized as active cells in subsequent analyses. Non-deconvolved traces were used for all downstream calcium imaging analyses. Cell spatial footprints were extracted (i.e., Figure 2) and aligned across imaging sessions using the MATLAB (R2020a, Mathworks) implementation of CellReg (Sheintuch et al., 2017). Multiday cell registration was completed based upon recommendations of CellReg probabilistic modeling. Alignment matrixes were manually evaluated using ROI visualization, and maximal distance shifts were adjusted if necessary to optimize cell tracking.

**Figure 2.**
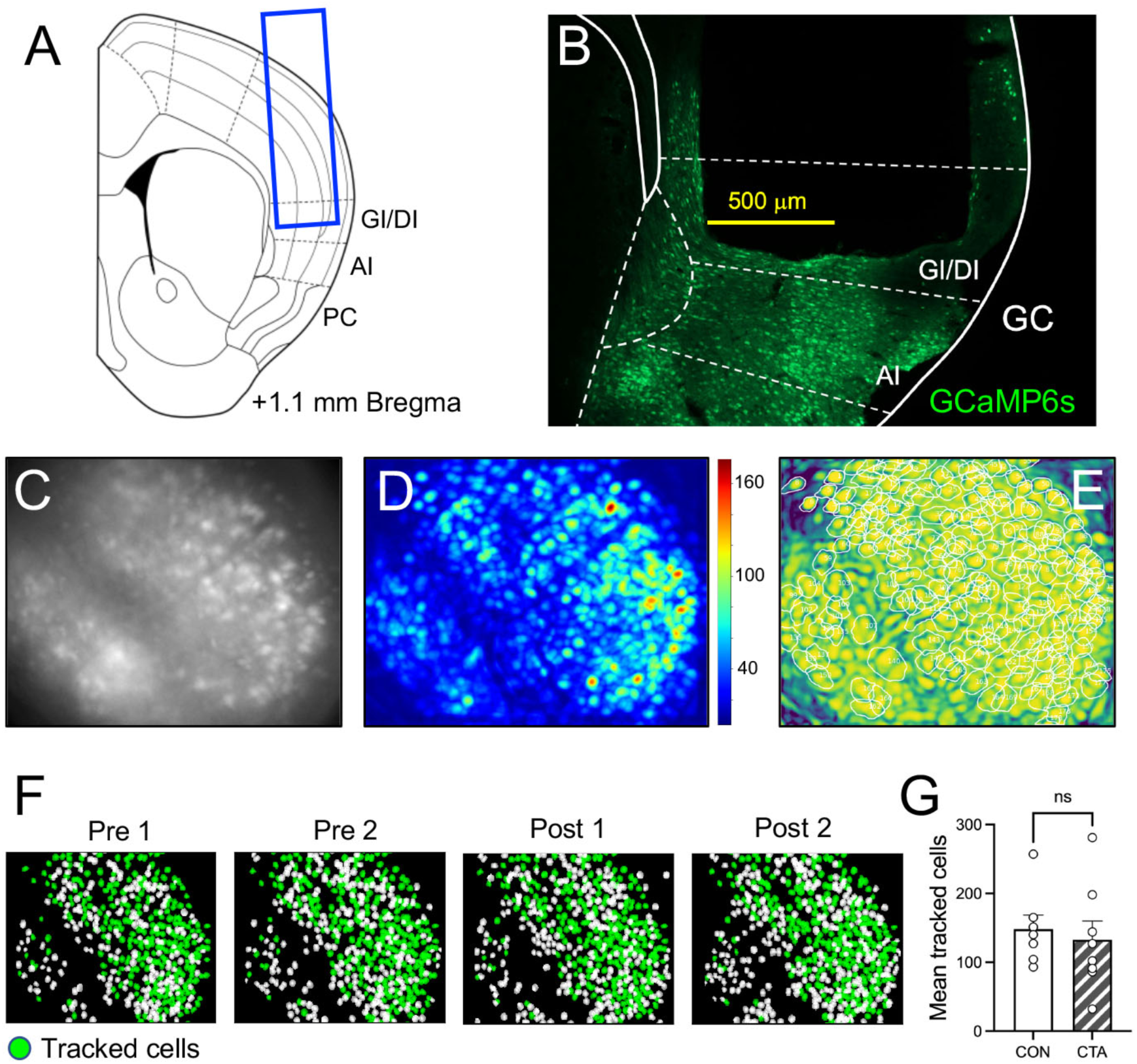
Calcium imaging procedure and analysis. A. Representative schematic of 1 mm-diameter GRIN lens placement in GI/DI (granular/dysgranular insular) cortex. B. Post-imaging fluorescent micrograph of cortical section showing GCaMP6 expression (green); the outline of GRIN lens implantation is discernable. C. Example of raw calcium fluorescent image in the miniscope field of view. D. Signal to noise overlay showing the degree of change in fluorescence through a session E. Spatial correlation overlay, with white circles indicating positively identified cells F. Spatial registration of cells over days, green indicating cells identified on all days. G. Average number of tracked cells (identified on all days) in control and CTA mice. AI, agranular insular cortex; PC, piriform cortex.

Further analysis of traces was conducted in R (Version 4.1.2) using custom scripts. Calcium traces were first aligned across days based on CellReg indices. Unless otherwise noted, cells were only included in analysis if they could be identified by CellReg in all experimental sessions. Taste-evoked change in fluorescence (ΔF) values were then calculated on a per-trial basis by subtracting the mean fluorescence evoked 5 seconds prior to licking from the maximum fluorescence evoked during the 5-sec licking period (Fletcher et al., 2017). As fluorescence data extracted from CaImAn has already been scaled by a background fluorescence value, ΔF, rather than ΔF/F, was calculated to describe neuronal responses (Giovannucci et al., 2019). For analyses in which cells are classified as excited or suppressed, significant responses were categorized as having at least a +/- 3.0 SD change in fluorescence (ΔF) compared to the 5-second pre-licking baseline period; otherwise, the delta values from all cells were used. Positive and negative delta values were considered in all analysis, excluding Entropy.

### Experimental Design and Analysis

Analysis was conducted on behavioral data, in most cases comparing 7 control with 8 CTA mice, and from the subset of 4 CTA mice undergoing extinction. Parametric statistical analyses (ANOVA or t-tests) were conducted using R or GraphPad Prism. Where appropriate (i.e., in the case of within factors), the Greenhouse-Geisser sphericity correction was applied. Post-hoc comparisons were made with the Bonferroni test. For imaging data, analysis was conducted on 2098 neurons (1061 cells identified in CTA animals, 1037 identified in controls; 448 cells retained from the CTA animals used in the extinction phase) recorded across all sessions and mice. We limited analysis to the set of cells identifiable on all days in the interest of understanding effects of learning on cell activity over time. Where multiple presentations of a taste stimulus were recorded in a single session, the responses for each stimulus were averaged, yielding one response value per cell and stimulus on a given day. Responsive cells from all mice were pooled by day. Entropy (H), a common measure of breadth of tuning used in taste recordings, was used to evaluate changes in cell tuning across days (Smith and Travers, 1979; Staszko et al., 2022).

Differences in mean taste-evoked responses were analyzed using one-way ANOVA and unpaired t-tests. Hierarchical clustering of lick counts and calcium responses was conducted using the Lance–Williams dissimilarity update formula with the complete linkage method. To compare changes in taste representations across days, we first performed multidimensional scaling (MDS; using pooled responses from mice) via principal coordinates analysis (Gower, 1966) to render stimuli in taste representational space, with the number of dimensions used in the solution selected via the scree method (Bieber and Smith, 1986). Subsequently, we calculated their proximity in taste space by comparing their Euclidean distances. The Euclidean distance (ED) between two representations for each day was calculated as:

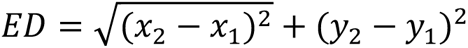

with x and y representing the coordinates of each taste in space and 1 and 2 indicating coordinates from two different tastes. Classification and discriminant analysis was conducted in Matlab with optimizable Support Vector Machines (Allwein et al., 2001; Furnkranz, 2002; Escalera et al., 2009, 2010).

### GRIN Lens Placement Verification

At the conclusion of imaging experiments, animals were anesthetized with ketamine/xylazine (100/10 mg/kg IP) and transcardially perfused using 4% paraformaldehyde (PFA). Following perfusion, the brain was further postfixed in PFA for seven days to improve fixation and delineation of the imaging window. Brains were then removed, cryoprotected, and sectioned in 40 um thick serial sections using a freezing microtome. Sections were mounted on slides and imaged using a Nikon Eclipse 90i fluorescent microscope (Nikon Instruments Inc., Melville, NY, USA) equipped with a digital camera and imaging software. A sample GRIN lens placement is shown in Figure 2A; all placements were plotted on schematic section diagrams. The anterior-posterior range of imaging sites was estimated at 0.75 to 1.5 mm anterior to Bregma, with most (9/15) cases at 1.0 – 1.1 mm. Although the 1.0 mm diameter of the lens allowed for coverage across most cortical layers, there was some variation among cases depending on factors such as the extent of GCaMP expression or angle of lens implantation. However, it was not feasible to identify cortical layer in the imaging lens field of view, so no analysis of this spatial parameter was attempted.

## Results

### CTA alters licking behavior

Prior to conditioning, all 15 mice received access to the taste panel for 2 days. Mean licks recorded per stimulus (groups combined; separate analyses showed that group assignment was not a significant factor) on the 2 test days prior to conditioning are shown in Figure 1B. Significant effects of both stimulus (F[6, 98] = 44.49, p < 0.0001) and day (F[1, 98] = 23.27, p < 0.0001) were found, as well as a significant stimulus x day interaction (F[6, 98] = 9.85, p < 0.0001, 2-way ANOVA). Essentially, licking of a number of stimuli increased from the first to the second pre-test, including increases for NaCl, KCl, and citric acid (ps < 0.05, Bonferroni tests). These effects likely indicate some degree of attenuation of neophobia (Arthurs et al., 2018), or are reflective of habituation to the stimuli. Still, it was apparent by day 2 that only one stimulus (high QHCl) provoked strong avoidance, with a mean lick count of 4.6 per 5 sec trial; all other stimuli were readily consumed, with mean lick counts > 21.0.

In the conditioning session, ingestion of LiCl by CTA mice depressed licking relative to the controls, who consumed equimolar NaCl. While both groups avidly and similarly consumed either stimulus in the first minute, the lick rate over the entire trial was higher for control mice drinking NaCl (Figure 1C; t[13] = 2.9, p = 0.012, unpaired t-test). All 7 control mice reached the 1000 lick limit in 5 minutes or less, while 5 out of 8 CTA mice did not reach 1000 licks in the 10-min time frame, indicating cessation of licking during the session. Although we did not quantify illness-related behavior during the test, these data are consistent with previous studies showing that ingestion of LiCl (but not NaCl) causes intake behavior to cease due to gastric malaise (Baird et al., 2005; Glatt et al., 2016).

For behavioral analysis of CTA effects, we focused on the comparison between the second pretest day and the (first) post-conditioning test day. Lick data to the panel of stimuli compared on pre- and post-tests for control and CTA mice in are shown in Figure 1D. CTA effects on licking were initially assessed with a 3-way mixed model ANOVA (group x day x stimulus). There was no main effect of either group or day, suggesting that for the most part behavior was equivalent across these factors. However, there was a main effect of stimulus (F[2.3, 29.85] = 96.68, p < 0.0001), and all interactions were significant. Based on these results, we conducted a series of two-way (group x day) ANOVAs for each stimulus. For NaCl, there were significant main effects of group (F[1,13] = 17.62, p = 0.001) and day (F[1,13] = 42.59, p < 0.0001) plus a significant interaction (F[1,13] = 33.29, p < 0.0001). Control animals’ licking of NaCl was essentially unchanged pre-post conditioning, but that of the experimental animals dropped precipitously, consistent with a CTA formed to LiCl that strongly generalized to NaCl (p < 0.0001, post-hoc test). The mean number of licks to NaCl in the post-test varied within the CTA group, but it is notable that lick number decreased from the pre-test value for every individual (Figure 1D). There were no significant group x day interactions for any of the other stimuli.

### CTA does not affect the number of taste-responsive cells in GC or response magnitude

Post-imaging examination of brains showed that GRIN lenses were located within GC, most commonly in granular or dysgranular insular cortex (GI/DI; example shown in Figure 2A-B). Examples of the imaging lens field of view, along with examples of fluorescence extraction, are shown in Figure 2C-E. We identified a subset of cells that could be detected on all days of the experiment (“tracked cells”; N = 139.9 ± 16.9 cells per animal; Figure 2F, G); this population of cells was subjected to further analysis.

Based on previous examinations of the effect of CTA in GC (Koh and Bernstein, 2005; Flores et al., 2018; Lavi et al., 2018), we anticipated there may be an increase in the number of neurons in GC responding significantly to the conditioned stimulus (either excited or suppressed by; for our purposes referred to as “perturbed”) following conditioning (pretest day 2 vs. post-test day 1 comparison). However, our analyses did not support this hypothesis. No significant effects on the percent of tracked cells responding to NaCl across group or day (pre vs. post) were found (Figure 3A; 2-way ANOVA). Moreover, expanding this assessment to include all taste stimuli also did not yield significant effects (3-way ANOVA). This lack of significant increase in percentage of responsive cells in the CTA group relative to controls was also true when considering excited and suppressed cells as subgroups (data not shown). Next, we evaluated the hypothesis that effects of CTA might manifest in GC neurons as an increase or change in response magnitude, especially to the conditioned stimulus, rather than a change in number of responsive neurons (e.g., Yamamoto et al., 1989; Moran and Katz, 2014). In terms of magnitude of excitation, we found no significant effect of group or day (or their interaction) on excited responses to NaCl (Figure 3C; 2-way ANOVA). When excited responses to all stimuli were combined, we found no significant effect of group or stimulus, but did measure an effect of day (F[1, 13] = 10.08, p = 0.007, 2-way ANOVA); the pooled excitation magnitude of all animals across stimuli increased significantly from the pretest to the post-test (t[14] = 3.3, p = 0.006, paired t-test). However, the group x day interaction was not significant. Assessment of suppression yielded no significant effects, either to all stimuli, or to NaCl only (Figure 3D; 2-way ANOVAs). Finally, we also tested for an effect of CTA on entropy of neurons to the 4 basic qualities of taste. Mean entropy for all tracked neurons on both pre- and post-days in either group was about 0.5; this intermediate value indicates fairly broad tuning, and is perhaps to be expected when considering such a large number of cells (Figure 3B). However, mean entropies did not change following CTA. A two-way ANOVA (day x condition) yielded no significant main effects of group or day, nor of their interaction.

**Figure 3.**
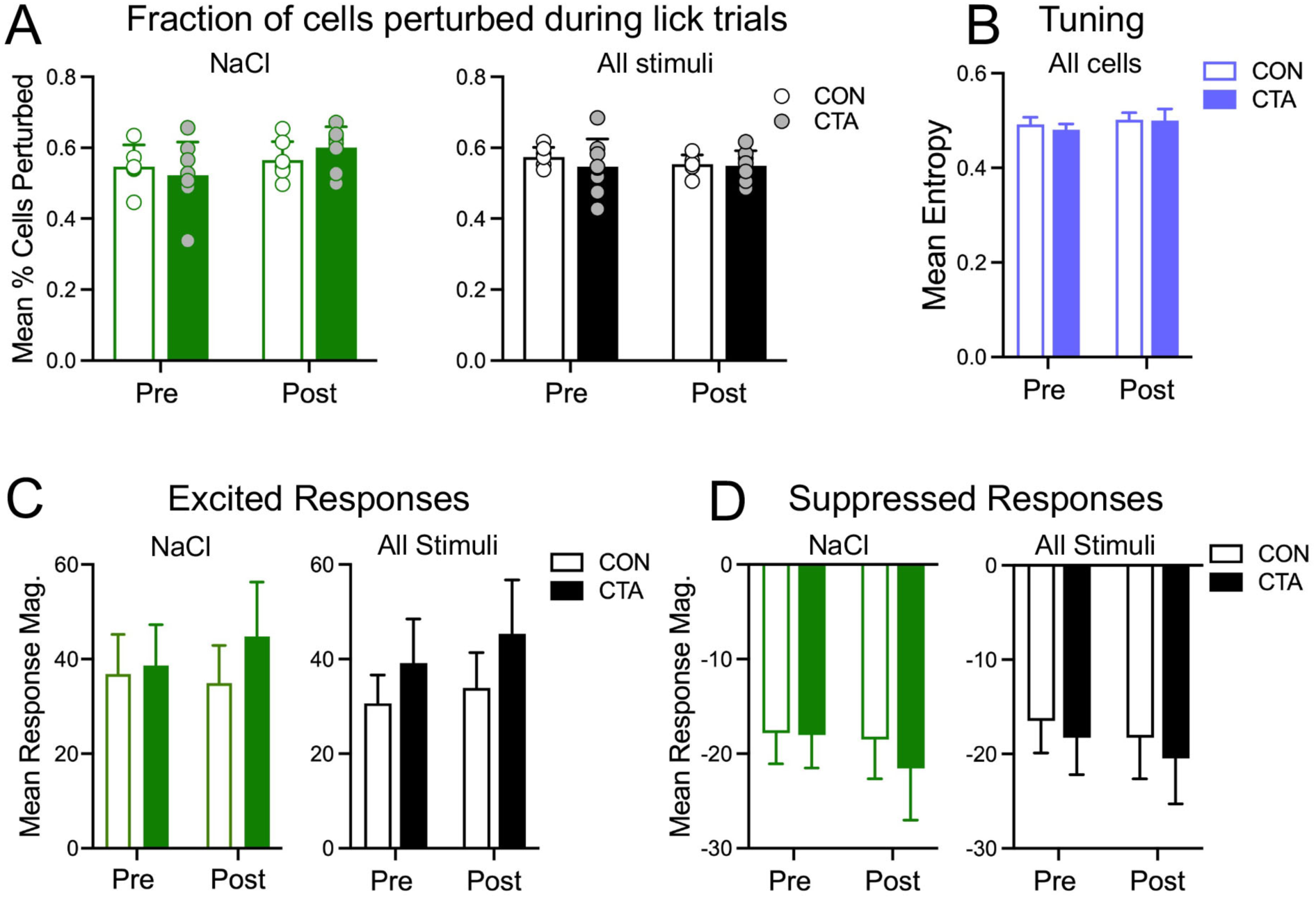
Basic activity measures pre- and post-CTA, averaged across animals and cells and compared between experimental groups. A. Fraction of cells perturbed (i.e., with significant excited or suppressed responses to taste stimuli) before and after conditioning in control (CON) and CTA animals, during trials with NaCl (green and white bars) or with all stimuli combined (black and white bars). Bars indicate mean values (± SEM) calculated across individual mice (circles). B. Breadth of tuning as measured by mean entropy across all cells, before and after CTA, in control and CTA mice. C. Average magnitude of significantly excited responses, either to NaCl alone (green and white bars) or averaged across all stimuli (black and white bars), before and after conditioning in control and CTA animals D. Average magnitude of significantly suppressed responses, either to NaCl alone (green and white bars) or averaged across all stimuli (black and white bars), before and after conditioning in control and CTA mice.

### CTA alters population coding of taste stimuli

To assess possible effects of CTA on taste coding, we compared the similarity of population representations of each taste stimulus between control and CTA groups. In taking this approach, we evaluated the hypothesis that NaCl might change its representation in terms of its similarity in neural space to other stimuli following learning (Moran and Katz, 2014; Staszko et al., 2022). The Euclidean distances between taste stimuli across identity and over time in the total neuronal population (data pooled across mice in each group) were calculated. Using the Scree method, we selected a two-dimensional solution to reduce the taste representations to their most prominent features. Because the dimension reduction was necessarily computed independently for each group, direct comparison between them is infeasible. Comparison within groups across days, though, was highly revealing (Figure 4A); we plotted multidimensional scaling (MDS) across 3 days, including both pre-test days and the post-test. In control mice, stimuli that were licked avidly cluster together, and their vectors across dimension 2 travel together in a tight cluster. On the other hand, high-concentration QHCl, which was avoided by mice, travels in the opposite direction, indicating evolving neural dissimilarity. In CTA mice, NaCl diverged from the cluster of appetitive taste stimuli on the post-test day, reflecting a shift in palatability following conditioning. These effects were tested statistically (Figure 4B): In two-tailed paired t-tests, mean distance between NaCl and appetitive taste stimuli was significantly greater post-CTA in the CTA animals (t[4] = 4.12, p = 0.015, paired t-test), but not in control animals. The post-test Euclidean distance (expressed in arbitrary units) between NaCl and high QHCl in the control group was 41.5, whereas in the CTA mice it was 19.7.

**Figure 4.**
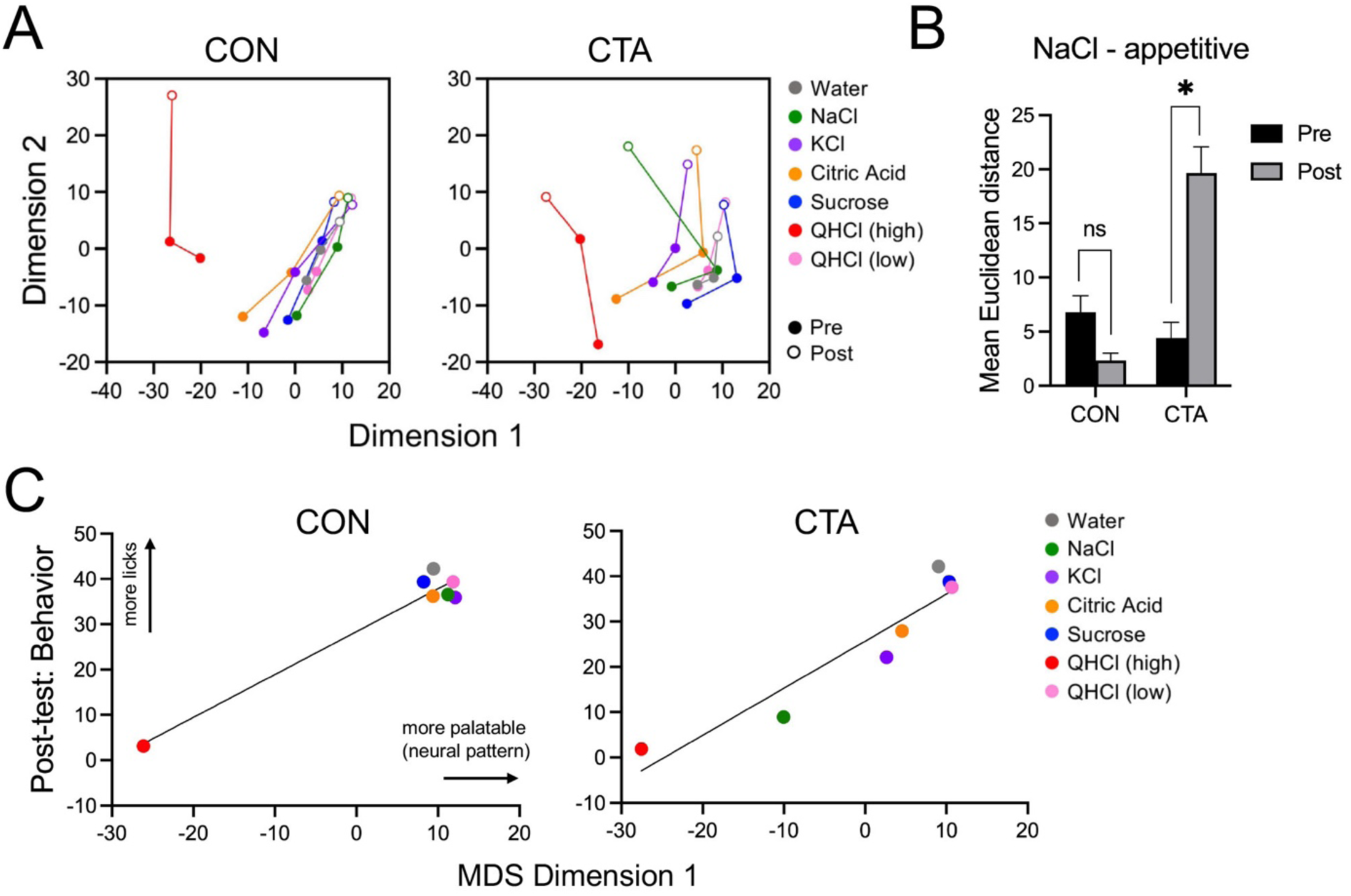
Relative taste representation in each experimental group as revealed by multi-dimensional scaling. A. Plot of the first 2 dimensions extrapolated by multidimensional scaling in control (CON) and CTA animals, showing the relationships among each stimulus within MDS space. B. Mean Euclidean distance between NaCl and other consumed (accepted) taste stimuli, before and after conditioning, in control and CTA animals. C. Plots demonstrating the high degree of correspondence between dimension 1 of the MDS and behavioral acceptance of each stimulus (i.e., mean licks) post-conditioning for control and CTA mice. Lines on plots indicate best-fit via simple linear regression of the data. Asterisks (panel B) indicate significance in post-hoc comparisons.

The tight relationship between the neuronal-derived MDS solution and the actual lick behavior, which are of course independently derived, is visualized in Figure 4C: Dimension 1 values and mean lick counts for each stimulus are highly correlated in either group (rs > 0.94; ps < 0.002). Furthermore, for CTA mice, the relationship between these two variables shows that following learning, NaCl has become less appetitive, reflected both in behavior and neural signature. Interestingly, minor effects of CTA on both KCl and citric acid (i.e., they have become slightly less appetitive) are also suggested by this data. This finding, while not immediately apparent in the behavior-only analysis (cf., Figure 1D), is consistent with a number of previous studies showing that a CTA to NaCl will generalize to some extent to some other salt and acid stimuli (e.g., Hill et al., 1990).

As an additional measure to corroborate the representational changes measured by MDS, we trained two Support Vector Machine (SVM) classification models, one per experimental group (Figure 5). We used SVM models to discriminate just the stimuli representing the four basic tastes (high QHCl, citric acid, NaCl, and sucrose) based on the activity evoked by those stimuli in the two pre-CTA sessions. We then applied those models to the data recorded on the day immediately following CTA, to assess how changing taste representations would impact the performance of the classifier. The SVM model trained and tested on the data recorded in control animals was able to correctly identify stimulus identity post-CTA with 75% accuracy; the model trained and tested on the data recorded in CTA animals performed at only 50% accuracy; this substantial reduction was due largely to the fact that the model misclassified both NaCl trials as the high concentration of QHCl (Figure 5A). Comparing the representation of NaCl recorded post-learning to a model constructed based on the pre-learning taste representations, it seems that the representation of NaCl post-CTA is more similar to the initial representation of QHCl than the initial representation of NaCl, an effect consistent with the findings of MDS.

**Figure 5.**
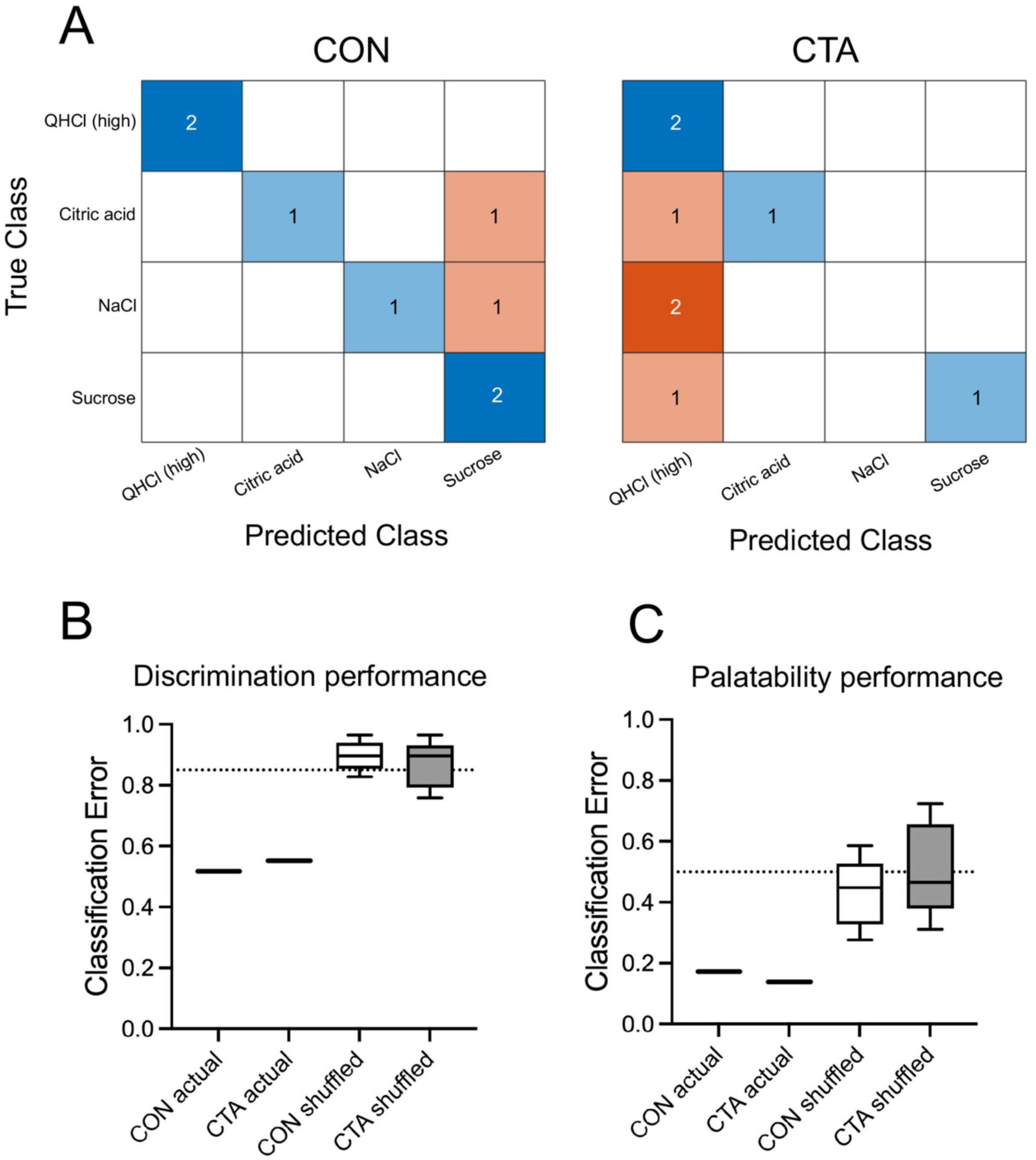
Classification models. A. Confusion matrices showing the predictions of a classifier trained on pre-conditioning data, which was asked to predict stimulus identity post-conditioning, in control (CON) and CTA groups. Blue squares show correct predictions; red squares, incorrect predictions. B-C. Boxplots of the cross-validation error rate of SVM classification learners, showing the prediction rate for either correctly labeled data or trials with shuffled labels; the dotted line shows the true chance rate for that prediction. Performance of classifiers in discriminating stimulus identity (B) and binary stimulus palatability (C) are shown in separate plots.

This apparent convergence of stimuli and failure of discrimination raised questions about the capacity of our GC imaging data to differentially encode various stimulus properties. In order to resolve those questions, we again turned to SVM classification, although the models were used differently for this task. By using leave-one-out cross-validation while fitting an SVM model on the entire data set, we were able to test the extent to which our data encoded stimulus identity and the hedonic nature of the stimulus as reflected by lick count. As an added control against over-fitting, we also benchmarked the performance of SVM prediction by feeding the algorithms versions of the data with shuffled labels. We first specified a model to predict stimulus identity. The classifier was trained with the evoked delta value of all cells per stimulus, averaged within each day. In a series of one-tailed one-sample t-tests (comparing the predictive power of a series of shuffled-label models either to chance or to true-label performance), we found that the classification error of shuffled SVM models was not less than chance (t[19] = 2.02, p = 0.971), and was significantly greater than the classification error of the real models (t[2.56] = 15.97, p = 0.0006). The true models for control and CTA animals had error rates of 0.517 and 0.552, respectively (Figure 5B). When asked to predict for each trial whether the average lick count of all animals in a given condition was higher or lower than the median lick count for all trials on that day (i.e., palatability), the shuffled models again failed to perform better than chance (t[19 = -1.03, p = 0.157; Figure 5C), and were themselves outperformed by the true models (t[10.98] = 9.05, p < 0.0001), which had error rates of 0.138 (CTA) and 0.172 (controls). Collectively, these analyses confirmed that taste-evoked activity in the population of tracked cells was able to encode information about both palatability and identity of taste stimuli, and this encoding was altered following CTA.

### Effects of extinction: reversal of CTA effects

Extinction of the CTA was first assessed by comparing licking of NaCl in control and CTA mice on post-test day 2 (Figure 6A). CTA mice had lower lick counts as compared with control mice throughout the initial 12 forced-drinking trials of NaCl (t[13] = 2.16, p = 0.049, unpaired t-test). However, when the unforced NaCl trial (i.e. presented in the panel of water and other taste stimuli) was considered, this difference was larger with the CTA mice showing greater aversion (t[13] = 4.37, p = 0.0008, unpaired t-test). Similar to our previous pre-post analyses, this difference in avoidance of NaCl between the two groups of mice did not correspond with a change in basic neuronal activity parameters in response to this stimulus. We found no significant group differences for percent responsive cells or in the mean excited or suppressed magnitude of response to NaCl for either forced or unforced trials) on post-test day 2 (data not shown). We next examined licking and imaging responses to NaCl in the subset of CTA mice (N = 4) who were tested for two additional days. For these analyses, only the unforced NaCl trials were considered, in order to examine the relationship among stimuli. In this group (Figure 6B), licking of NaCl changed significantly over days (F[2.2, 6.6] = 6.43, p = 0.026, 1-way ANOVA). It is evident that while consumption of NaCl is lower on post-test days 1-2 relative to the pretest baseline, it rebounds by post-test day 3, suggesting extinction of the CTA.

**Figure 6:**
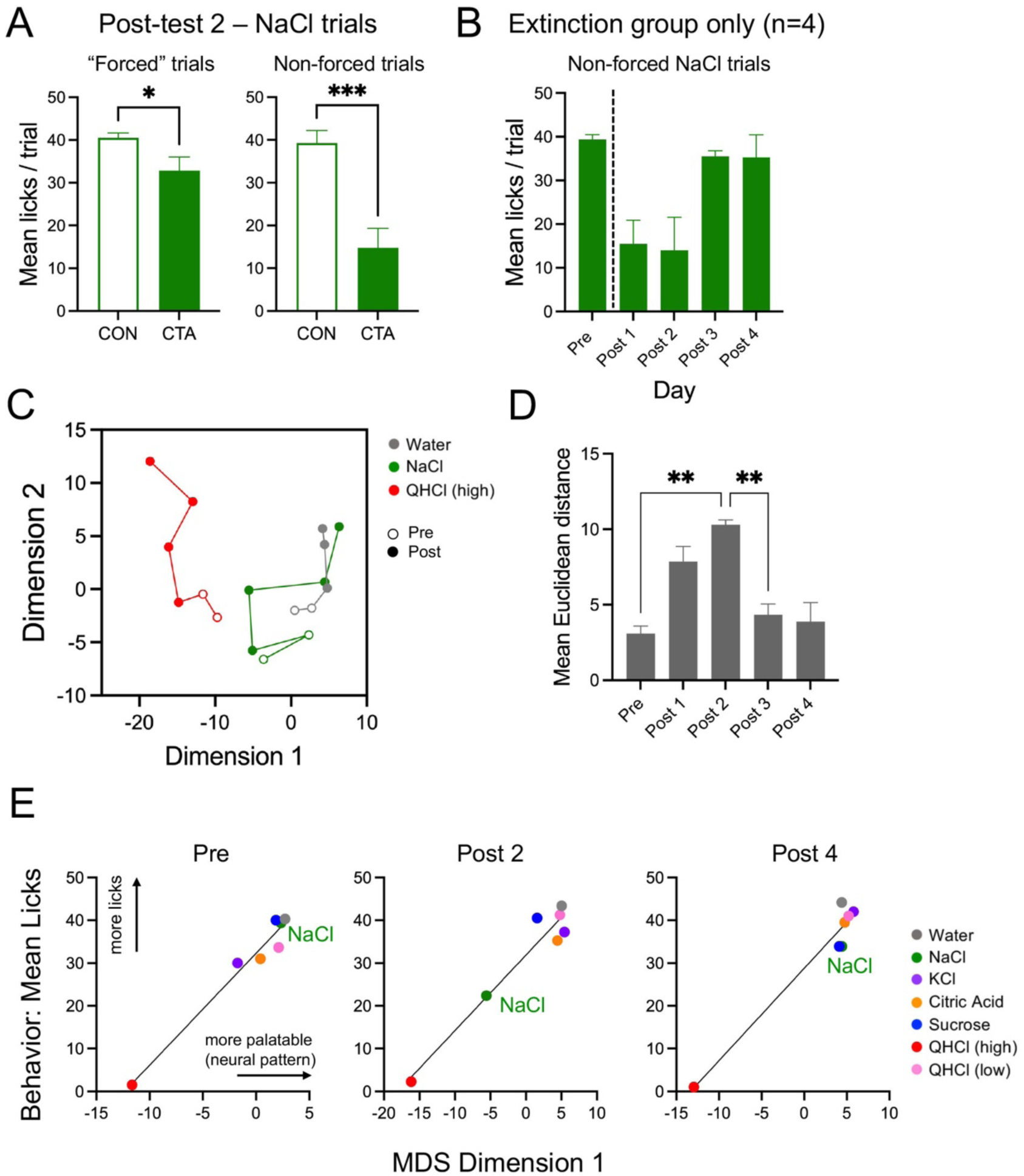
Analysis of extinction. A. Mean licks per trial for NaCl in control (CON) and CTA animals post-conditioning, separated into 12 consecutive “forced” NaCl trials (left), or the drinking of NaCl embedded in the randomized panel (non-forced; right). B-E: Data is shown for extinction-only mice. B. Mean licks per trial of NaCl by day. C. Plot of the 2 dimensions of MDS over the full course of testing, showing only selected stimuli (water, NaCl, high QHCl) for legibility. D. Mean Euclidean distance between NaCl and all behaviorally accepted stimuli over pre-test 2 and all post-test days. E. Relationship between MDS dimension 1 and behavioral acceptance of all stimuli over CTA and extinction; data is plotted for the pre-test, and post-test days 2 & 4. Lines on plots indicate best-fit via simple linear regression of the data. Asterisks (panels A, D) indicate significance in t-tests or post-hoc comparison tests.

We next examined the effects of extinction on population coding with multidimensional scaling in the extinction-only group (Figure 6C). Here, all pre- and post-test days are plotted for QHCl (high), NaCl, and water. Following conditioning, the NaCl vector initially moves away from water and towards QHCl, but by post-test days 3-4, it overlaps with water, reflecting similarity in representation between these stimuli and extinction of the CTA. A comparison of the mean Euclidean distance between NaCl and tastes other than high-concentration QHCl indicated a significant effect of day (F[4, 16] = 13.14, p = 0.003, 1-way ANOVA). Distance peaked on post-test day 2; mean Euclidean distance was significantly different between this day and both the pre-test and post-test day 3 (ps < 0.016, post-hoc tests). When MDS dimension 1 (palatability) was plotted against lick behavior for mice in this group, a strong relationship between these variables was evident with NaCl moving towards high QHCl along the regression line following conditioning, and then back towards the other appetitive stimuli by the final post-test day (Figure 6E).

## Discussion

In this study, we used miniscope imaging in behaving mice in order to understand changes taking place in a stable population of cortical neurons following CTA learning and extinction of that learning. Understanding the manner in which taste representations in GC may altered by CTA is crucial to revealing the function of GC itself. Based on previous research, we hypothesized that conditioning may change these representations in two ways: First, the identity of the CS may become more salient and singular, facilitating more efficient identification. For example, there is an increase in the number of CS-responsive neurons in GC (in rats) following CTA relative to controls in studies measuring neural activity with cFos (Koh and Bernstein, 2005; Uematsu et al., 2015; Flores et al., 2018). In our study, we did not observe a similar increase in neuronal activity in response to the CS (NaCl) following CTA, either in the percentage of cells activated or magnitude of responses. However, evoked physiological activity and Fos expression is not always correlated in a population of neurons, and expression generally indicates excitation, not inhibition or relative strength of response (Fenelon et al., 1993; Lara Aparicio et al., 2022). Previous physiological or imaging studies demonstrated that changes in response were limited to subsets of taste-responsive neurons (Yamamoto et al., 1989; Yasoshima and Yamamoto, 1998; Lavi et al., 2018). For example, a recent 2-photon imaging study found an increase in number of responsive cells and mean excited response following CTA in GC cells that project to the basolateral amygdala, but not in non-projection cells (Lavi et al., 2018). On the other hand, the results of several ensemble recording studies casts reasonable doubt on whether changes in activity in individual cortical neurons are correlated with the behavioral changes seen in response to the CS following CTA learning or extinction (Moran and Katz, 2014; Arieli et al., 2022). In these studies, single-neuron responses to the CS (sucrose or saccharin) changed following CTA, both in terms of response magnitude (either increase or decrease) and time course, although these changes averaged out across the population. Moreover, the changes in cell response did not revert following CTA extinction, suggesting this parameter did not play a role in representation of behavior.

In the current study, we found effects of CTA on the representational valence of NaCl that correlated with a change in behavior following learning. These effects were discernable at the population, rather than single-cell level. When the MDS representations of tastes were examined in control mice, NaCl clustered with other consumed stimuli, forming a distinct difference from the lone behaviorally-avoided compound, high-concentration quinine. This effect was mirrored in experimental animals prior to CTA. However, after learning an aversion, the representation of NaCl began moved towards that of high QHCl, with a significant increase in Euclidian distance from consumed stimuli. Notably, however, while the representation of NaCl closed much of its initial distance from QHCl after CTA, the two aversive representations never achieved proximity comparable to the distance between behaviorally accepted compounds, a phenomenon that reflects previous findings (Moran and Katz, 2014). This suggests that although NaCl has become aversive following learning, it does not share taste identity with this stimulus – notably, the SVM analysis suggests that although there may some confusion between the two, responses to all stimuli in both CTA and control mice support a degree of stimulus discrimination. The changes in palatability of NaCl we observed were reversed by behavioral extinction of the conditioned aversion, also in accordance with some previous studies (Accolla and Carleton, 2008; Moran and Katz, 2014).

It is striking how the neuronal population response in GC to taste stimuli accurately reflected the behavior, with neural patterns correlated among stimuli that mice licked avidly, and distinct from those they avoided. Stimuli can be distinguished by their quality (i.e., their distinct chemical nature), which is sometimes casually conflated with certain assumptions of palatability, derived mainly from human perception and extrapolated from the orofacial responses of animals (e.g., Grill and Norgren, 1978; King, 2018; Dolensek et al., 2020). However, as mentioned earlier, behavioral acceptance of a given stimuli is highly subject to context, and at times may run contrary to “liking”, as in a state of physiological need when an animal may be willing to consume a taste stimulus it would otherwise avoid (Scalera, 2000; Mast et al., 2017; Miranda et al., 2023). We found that in our observations, the behavioral response to a typical taste stimulus set fell into a more or less binary pattern – licked or not licked—which could not be predicted strictly by stimulus quality. For example, mice in our experiment consumed 0.3 mM (low concentration) QHCl robustly, whereas they strongly avoided 0.01 M QHCl. In previous licking studies, 0.3 mM provoked moderate to low avoidance in thirsty mice, based on the strain used (Glendinning et al., 2001; St. John and Boughter, 2004; Boughter et al., 2005); avoidance was much greater in 2-bottle tests with unrestricted mice (Bachmanov et al., 1996). In other words, mice may readily drink a weaker concentration of a bitter stimulus when motivated by thirst. These two concentrations of QHCl, which share their taste quality and under some circumstances can elicit similar behavioral effects, were consumed differently in our experiment and so evoked different representations in GC. By contrast, 0.3mM QHCl and 0.5M sucrose, though dissimilar in quality, were consumed similarly in our paradigm, and elicited much more similar cortical representations. This trend holds true even with a single concentration of stimulus: A CTA generalizing to NaCl alters the behavioral response to, but not qualitative nature of, of that stimulus. This learning was commensurate with an alteration of the cortical representation of NaCl stimulus from appetitive to aversive.

Our results are consistent with the idea that the gustatory representation in GC plays a predominant role in palatability, or perhaps “consummability”, of taste stimuli present in foods and fluids. This does not mean that stimulus quality is not encoded in the activity of GC neurons: Cortical recording or imaging studies in mice demonstrate encoding of both stimulus identity and palatability (Fletcher et al., 2017; Levitan et al., 2019; Bouaichi and Vincis, 2020; Chen et al., 2021). In the time-epoch taste coding model established by Katz on the basis of GC ensemble recordings (e.g., Katz et al., 2001; Levitan et al., 2019), the stimulus detection/discrimination phase are brief (> 1s) and quickly give rise to a longer-lasting palatability signature. In other words, appetitive vs. aversive becomes the dominant cortical representation. Here, by combining imaging of the same population of GC neurons day after day in mice actively tasting and licking stimuli, we provide evidence that this representation is not best described by the predicted hedonic attributes of the stimulus, but by the actual behavior the animal displays. In our paradigm, lick behavior is modulated by thirst – recent work suggests that neural activity patterns in part of insular cortex (in an area slightly more posterior than the current study) are in fact linked to physiological need states, including hunger and thirst (Livneh et al., 2020).

The implication of an organization of taste representations that is heavily dominated by palatability is a profound association of GC with ingestive choice, although the direction of this relationship is not immediately clear. It is perhaps an easy assumption to make (but not necessarily the correct one) that this ingestive choice behavior originates in GC. However, there remains the possibility that the cortical taste representations are driven by the sensory consequences of the behavior, rather than driving the behavior itself. To some extent this is inevitable; the sensory inputs to GC dictate some percentage of its activity (GC receives taste-related inputs from multiple areas, including from the brainstem, gustatory thalamus, and basolateral amygdala), and there is evidence that an intact GC may be unnecessary for at least the most basic judgement of palatability (Grill and Norgren, 1978; Braun et al., 1982; King et al., 2015). However, the anatomical evidence of GC outputs, including robust projections to motor and motor-related areas (Gehrlach et al., 2020), preclude the exclusive dominance of sensory representation in this brain region. Awake recording studies in rats demonstrate close temporal correspondence and a degree of dependency between the cortical taste response and orofacial behavior (Li et al., 2016; Sadacca et al., 2016). Furthermore, there is ample evidence that disruption of the GC disrupts many aspects of taste-related executive function beyond the most basic (Sastre and Reilly, 2006; Blonde et al., 2015; Lin et al., 2015; Schiff et al., 2018; Vincis et al., 2020; Jung et al., 2022). Given the additional consideration that the preponderance of sensory input into GC might be assumed to be taste-rather than somatically evoked, and that the activity we record in GC does not co-vary especially well with strict taste quality, it follows that the tight correspondence of GC representations with behavior is a causative element, rather than caused. As such, it is our conclusion that the representations of tastes in GC are dominated not by discrimination of identity, but rather the decision of imminent action.

